# Development of a high throughput continuous fluorescence anisotropy-based assay to analyse peptidoglycan-digesting enzymes

**DOI:** 10.64898/2026.10.19.760132

**Authors:** Stefano Camaione, Charis M. Thompson, Nicholas S. Briggs, David I. Roper

## Abstract

Bacterial cell wall processing enzymes have a central role in the control of the construction, remodelling and breakdown of the peptidoglycan wall that protects bacteria from osmotic rupture. Many of these enzymes have biologically essential functions and exert mechanistically diverse activities on a complex polymer substrate. Despite this biological importance they are relatively understudied and there are significant technical challenges in measuring the activity of individual enzymes against the complex chemical structure of the bacterial cell wall peptidoglycan. To generate a suitable substrate, bacterial cells are grown with a fluorescent D-alanine analogue, allowing the isolation of natively fluorescently labelled PG. This material is then used to measure interactions with PG-targeting enzymes and regulatory proteins. The resulting assay enables analysis of multiple enzyme–substrate combinations within minutes while retaining molecular specificity. This approach provides a robust, cost-effective alternative to conventional PG activity assays and could accelerate the identification and characterisation of novel antimicrobial compounds targeting bacterial cell wall metabolism. Our results indicate this assay has the potential to be employed in high-throughput inhibitor discovery, making it effective for a broad-spectrum assay for bacterial peptidoglycan research applications.

## INTRODUCTION

The peptidoglycan (PG) layer is an intricate polymer that encapsulates the inner (or cytoplasmic) membrane of most bacteria. It is involved in maintaining the species-specific cell shape, as well as functioning as a scaffolding structure for the anchoring of proteins; while being flexible and porous enough to ensure secretion of quorum sensing signals and virulence factors [1,2]. The PG also plays a fundamental role in counteracting the internal turgor pressure of the cell itself, preventing it from rupturing [3]. Based on physical differences in the cell wall architecture, bacteria are classified as Gram positives, presenting a thicker PG layer and a single membrane delimiting the cytoplasmic space; and Gram negatives, presenting a thin PG layer enclosed in the periplasmic space between an inner and an outer membrane [4]. Outside of this classification are organisms of interest like Mycobacteriaceae, which present modifications on the glycan backbone of the PG, alongside an arabinogalactan layer and an outer membrane decorated with mycolic acids [5], and Chlamydiaceae, for which PG existence has only recently been proven [6].

As the name suggests, the PG is a polymer composed of a glycan backbone formed by repeating units of two sugars, *N*-acetylglucosamine (GlcNAc) and *N*-acetylmuramic acid (MurNAc) that are linked by a ß-1,4 glycosidic bond [1]. The MurNAc sugar is then linked via amide bond to an _L_Ala residue, which represents the N-terminal portion of the peptide stem of the PG (Fig. 1). The peptide stem fundamental structure comprises an N-terminal L-Ala, followed by a D-Glu, an mDAP (L-Lys in Gram positives [7]) and a D-Ala-D-Ala dipeptide at the C-term. The D-Glu in position 2 is amidated in certain species to *i*d-Gln [2]. The peptide stem itself is interconnected by a complex net of crosslinks between the mDAP in position 3 [4] (_L_Lys in most Gram positives) and the _D_Ala in position 4, formed by D,D-transpeptidases (3-4 crosslinks). Some bacterial species rely instead on the combined action of D,D-carboxypeptidases, cleaving the C-terminal _D_Ala [8], and L,D-transpeptidases, which will form 3-3 crosslinks [9].

**Figure 1:**
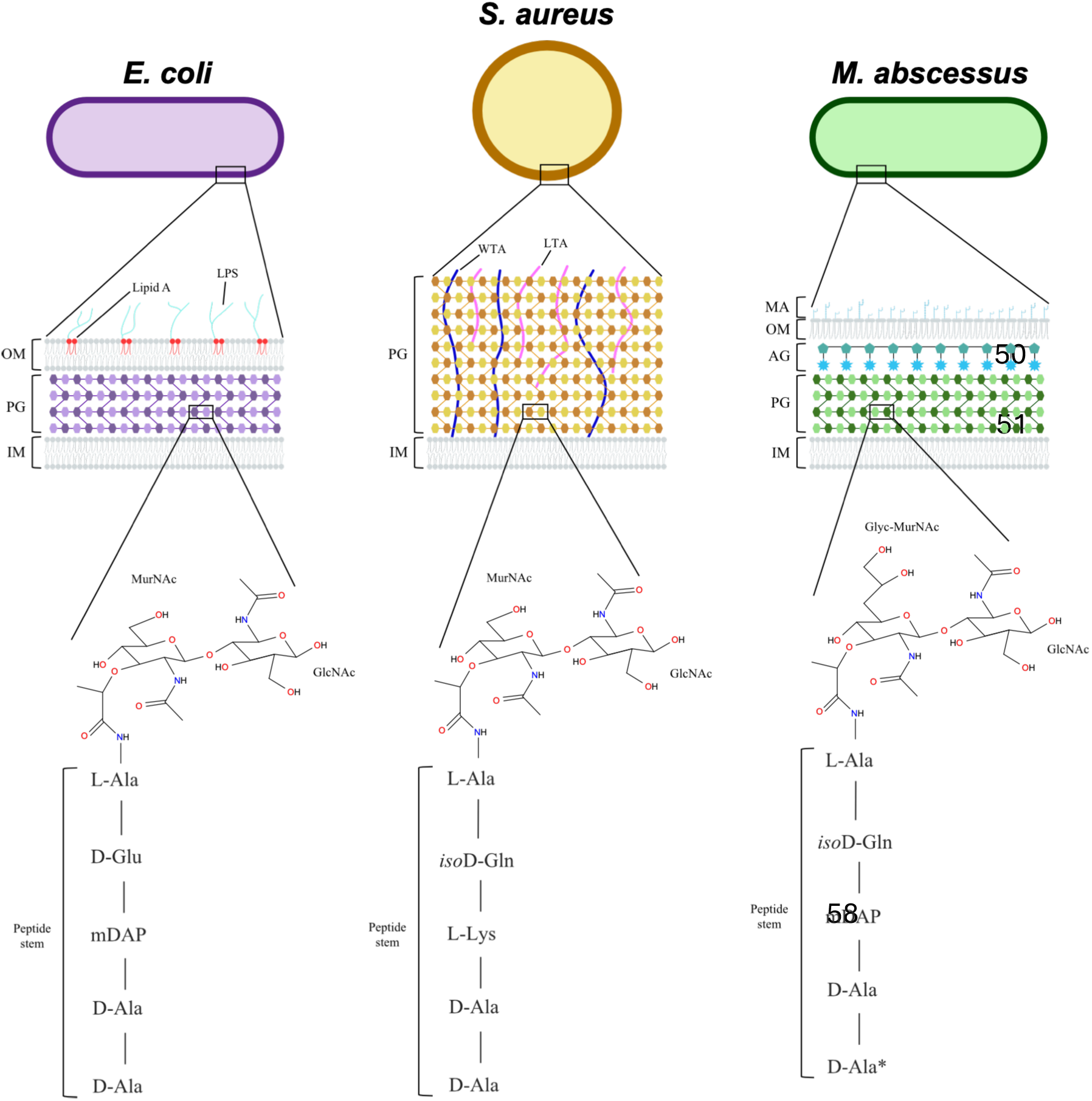
schematic representation of Gram negative (purple, *E. coli*), Gram positive (yellow, *S. pneumoniae*, *S. aureus*) and mycobacterial (green, *M. abscessus*) peptidoglycan, highlighting the differences in PG layer thickness and composition. The peptide stem is not representative of the presence of crosslinks and peptide bridges for ease of reading. *: the C-terminal D-Ala in *M. abscessus* is normally not present in the mature PG, as it gets cleaved by D,D-carboxypeptidases in 3-3 crosslinked cell walls. OM = outer membrane; PG = peptidoglycan; IM = inner membrane; LPS = lipopolysaccharide; WTA = wall teichoic acid; LTA = Lipoteichoic acid; MA = mycolic acid; AG = arabinogalactan; GlcNAc = *N*-acetylglucosamine; MurNAc = *N*-acetyl muramic acid; Glyc-GlcNAc = glycolylated MurNAc

The PG architecture is maintained by a delicate equilibrium of synthesis and degradation. For a more in-depth description of these mechanisms, we encourage the consultation of recent reviews on the subject [10, 11].

Enzymes involved in bacterial cell wall biosynthesis have traditionally been targets for drug discovery research, with penicillin binding protein class enzymes (PBPs), being among the most studied targets [12, 13]. Often overlooked in this context are the enzymes responsible for PG maturational turnover, where there is considerable apparent redundance of function. This implicates an important biological necessity, made difficult to analyse due to the lack of analytical techniques to easily assay individual function. PG-digesting enzymes are involved in the controlled degradation of the PG layer required for cell wall remodelling [14, 15] and may therefore represent promising targets for novel antimicrobial drug discovery [16]. Pharmacological disruption of these tightly regulated processes could dysregulate bacterial cell-wall metabolism and provide a new strategy for addressing antimicrobial resistance (AMR), a major global health threat that is predicted to surpass cancer as a leading cause of death [17].

When studying an enzyme as a possible drug target, it is important to ensure accurate enzymatic activity and, more importantly, accurate inhibition by a candidate compound on a defined substrate. To this end, most enzymatic activity assays rely on reaction product formation, substrate degradation rate, enzyme-substrate interaction, or formation of an intermediate complex [18], to ensure calculation of enzyme kinetics and understand inhibition mechanism. In this context, PG represents a difficult substrate to work with, often due to the high specificity of PG hydrolases [19, 20], which restricts the choice of activity assays to laborious spectrophotometry-based measurements [21, 22, 23] or cost-demanding and highly technical HPLC- and UPLC-MS separation methods [24, 25]. While these methods are accurate and allow for inhibition confirmation of selected compounds, they often rely on primary screens to reach through high throughput (HT), thus elongating the drug discovery timeline and increasing overall cost.

In this work, we propose the repurposed use of a known technique, fluorescence anisotropy (FA), to analyse native PG both as a substrate for PG hydrolases and as an interactor for regulatory PG-binding proteins. FA is based on the selective excitation of fluorophores by polarised light, thus allowing for the discrimination of fluorescent molecules based on their size [26]. This principle is already widely used for the study of protein-protein [27] and protein-small molecule [28] interactions, as well as to analyse enzymatic activity [29]. FA as a technique is already a recognised and established method that is approved by regulatory agencies as a drug discovery tool [26]. Based on this, we modified existing PG isolation methods to obtain native fluorescently labelled PG by growing cells in the presence of a fluorescent D-Ala supplement. The extracted fluorescent PG was used as a substrate for known PG-targeting enzymes, thus developing an assay capable of analysing different enzyme-substrate combinations in a matter of minutes. This robust system represents an attractive alternative to traditional screening tools, decreasing sample preparation time whilst retaining protein-ligand specificity.

## METHODS

### Enzymes and bacterial culture conditions

LytA and MltD_LysM_ were expressed and purified in-house (see below). Mutanolysin was purchased from Sigma Aldrich, lysozyme was purchased from Invitrogen, lysostaphin was purchased from Merck.

Bacterial species used in this study, alongside the relative culture conditions, are enlisted in table 1.

**Table 1:**
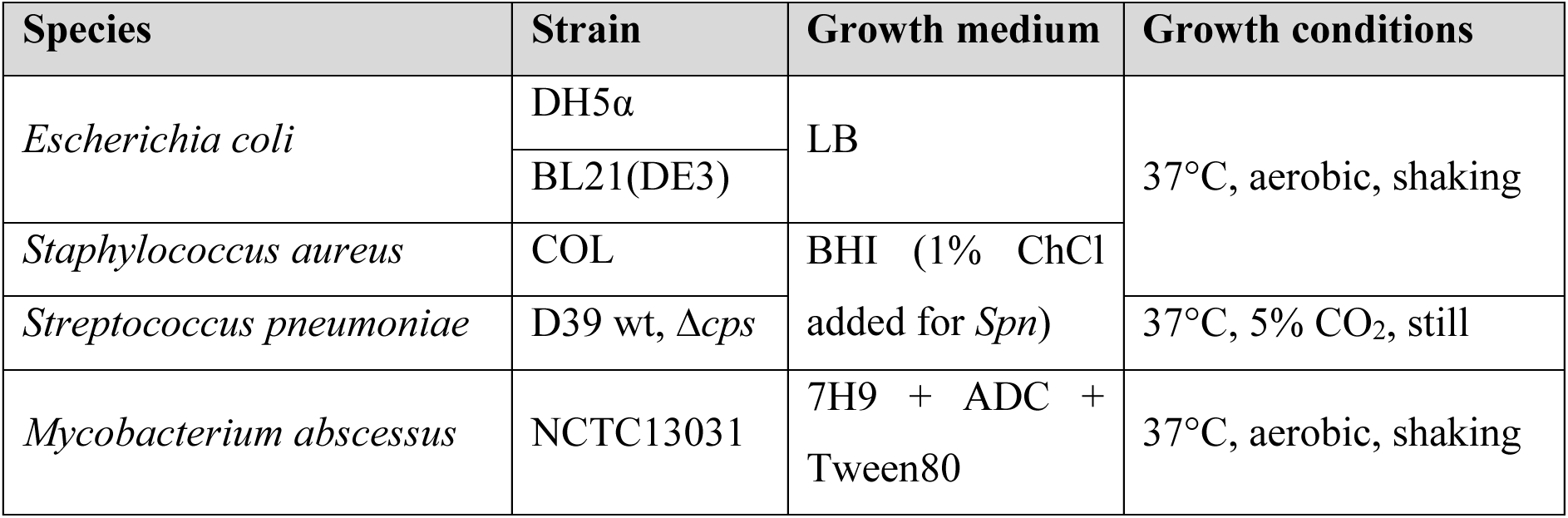
bacterial species and relative culture conditions.

### Construct cloning and protein purification

The *lytA* gene was amplified from *S. pneumoniae* D39 genome using primers LT1 and LT2 (Table S1), cloned into a pET-28b(+) vector via Gibson assembly and confirmed by Sanger sequencing.

*E. coli* BL21(DE3) expressing the protein of interest was grown overnight shaking (180 rpm) at 37°C in Luria Broth (Merck) and subsequently diluted 1:20 in fresh 2YT (16g/L tryptone, 10 g/L yeast extract, 5 g/L NaCl).

To obtain LytA, cells were grown at 37°C with vigorous shaking (220 rpm) until exponential phase was reached (OD_600_ 0.4-0.6), at which point protein overexpression was induced by adding isopropyl ß-D-1-thiogalactopyranoside (IPTG, Invitrogen) to a final 1 mM concentration. Temperature was dropped to 28°C and induction continued for 4 hours. Cells were harvested by centrifugation (6891*g* for 10 min), resuspended in lysis buffer (20 mM Tris-HCl pH 7.5, 500 mM NaCl, 10% glycerol) and lysed via sonication. Soluble proteins were separated via centrifugation (50000 *g* for 30 min) and loaded onto 1 ml of Ni-NTA resin (Thermo Scientific). Resin was washed with lysis buffer, and protein elution was achieved by washing with lysis buffer supplemented with 300 mM imidazole. Fractions containing LytA were pooled and buffer exchanged into 20 mM Tris-HCl pH 8, 20 mM NaCl, 10% glycerol before being loaded onto a 5 ml HiTrap Capto Q column (Cytiva) to be further purified via anion exchange chromatography. Column was washed with loading buffer and LytA was eluted with a 20 CV gradient to 1 M NaCl. Protein purity was assessed via SDS-PAGE (Fig. S1) and LytA was finally dialysed in storage buffer as described by Mellroth *et al*. [30] (20 mM Tris-HCl pH 7.5, 200 mM NaCl, 5 mM ChCl_2_, 10 µM ZnCl_2_).

### Peptidoglycan extraction

To extract peptidoglycan, we adapted the methods previously published by Kühner *et al.* [24] and Mahapatra *et al.* [31].

To maximise labelling and PG recovering, cells were grown overnight (or for 72 hours in the case of *M. abscessus*) at 37°C shaking (180 rpm) in 10 ml of the appropriate medium (50 ml in CO_2_ atmosphere with no shaking for *S. pneumoniae*) in the presence of 5 µM OGDA (Tocris) to label the PG. Cells were harvested via centrifugation (3220*g* for 10 min), resuspended in 50 mM Tris-HCl pH 7.5 + 1 M NaCl (*E. coli* and *S. pneumoniae*) or 1% SDS (*S. aureus* and *M. abscessus*) to a final cell concentration of 10 OD/ml and boiled at 100°C for 25 min. Cells were washed three (NaCl treatment) or six times (SDS treatment) with milliQ water. SDS removal was checked via chloroform/methylene blue quantification [32] and the resuspended cells were incubated in a sonication bath for 30 min. Peptidoglycan-associated proteins were digested and removed by incubating the sonicated cells for 2 hours at 37°C with 500 µg/ml trypsin in 2 mM MgCl_2_ (final concentrations). Upon digestion, the samples were boiled at 95°C for 5 min and the cells resuspended in 1 ml milliQ water. *E. coli* PG was ready to be used at this point. *S. pneumoniae* and *S. aureus* PG was treated for 4 hours at 37°C with 1 N HCl to remove wall teichoic acids. For the extraction of *M. abscessus* PG, the trypsin-digested samples were suspended in 89.1 mM KOH in methanol and incubated at 37°C shaking for 4 days to remove mycolic acids. The pellet was then washed twice in MeOH, and the resulting arabinogalactan-PG was resuspended in 0.05 N H_2_SO_4_ and incubated at 37°C for a further 5 days to remove arabinogalactan. The resulting PG was washed with mQ water until pH was between 5 and 6 and finally resuspended in phosphate buffer (12.5 mM NaH_2_PO_4_, 5 mM Na_2_HPO_4_ pH 6.2).

Final yield was calculated by pre-weighing the final storage vial, then weighing it again before resuspending the pellet in storage buffer. Given the non-solubility in water of non-digested peptidoglycan, it is critical to thoroughly mix the suspension by vortexing or vigorous shaking before pipetting in the wells to use as substrate; failure to do so will result in non-consistent repeats and loss in analytical power.

Samples of each main passage were taken and analysed for general fluorescence to optimise the procedure.

Fluorescent D-Ala retainment in the extracted PG was assessed by measuring fluorescence intensity in a CLARIOstar Plus plate reader (BMG Labtech).

Our method has proved to be scalable to several litres of starting culture with no effect on the final yield of fluorescent substrate (Fig. S2).

### Fluorescence anisotropy

All fluorescence anisotropy measurements were carried out in a CLARIOstar Plus plate reader (BMG Labtech) using an excitation wavelength of 482-16 nm and an emission wavelength of 530-40 nm, with dichroic lens set at 504 nm. Incubator temperature for the reader was set at 37°C. Due to difference in general fluorescence between the different species, focal height was assessed before the start of each experiment, and gain was set at 95% maximum intensity to increase sensitivity. Cycle time was determined based on the predicted minimum time to complete a full layout read, so to have the minimum time delay between readings on the same well.

To measure enzyme activity, substrates were resuspended in phosphate buffer at 50 µg/ml final concentration, then 9 µl were transferred to the wells of a pre-warmed black 384 low-volume plate (Greiner). Immediately before reading, 1 µl of enzyme solution (pre-diluted in phosphate buffer to obtain a final concentration of 200 µg/ml) was added to the wells and the plate was read for 45 min.

To measure enzyme kinetics, serial 3-fold dilutions of substrate were prepared. To have better control over the sample, the dilutions were prepared in 96-well plates (U-bottom, Greiner) in 45 µl total volume, then 9 µl of each dilution was transferred to the assay plate and the enzyme was added as described. Initial velocity was calculated from the slope of the first 2 min after blank subtraction and plotted against substrate concentration to obtain K_m_.

IC_50_ was measured for *N*-bromosuccinimide (NBS) against mutanolysin by incubating the enzyme with serial 3-fold dilution of NBS before adding them to the 50 µg/ml substrate as described. Initial velocity was again calculated as described and plotted against inhibitor concentration.

### Flow Induced Dispersion Analysis

Flow Induced Dispersion Analysis (FIDA) was employed to analyse the digested labelled muropeptides as a way to both assess retainment of fluorescence and characterise the molecules based on their absolute hydrodynamic ratio. Soluble muropeptides were pushed into a permanently coated capillary (FIDAbio, Denmark) at 1200 mbar for 60 s, following a pulse of phosphate buffer at 3500 mbar for 25 s to fill the capillary. Taylor conditions were calculated for an expected hydrodynamic ratio < 4 nm via FIDA analysis software. Molecules passing was monitored by a 480 nm fluorescence detector, and the resulting taylorgrams were analysed applying the Stokes-Einstein equation. All measurements were conducted on a FIDAneo (FIDAbio).

## RESULTS

### Extracted labelled peptidoglycan can be used as substrate in a fluorescence anisotropy assay

PG from *E. coli*, *S. pneumoniae*, *S. aureus* and *M. abscessus*, representing different chemical forms from diverse species, was labelled with fluorescent D-Ala (OGDA) by adding the dye to the starter culture and incubating to late stationary phase, ensuring maximum saturation of the PG and retention of the highest level of fluorescence upon extraction.

Testing of the extracted labelled PG was performed against non-labelled PG of the same species, showing retainment of the fluorescent dye. This, to our knowledge, is the first example of a natively labelled PG being extracted from cells, without the requirement of post-extraction modifications [33].

To streamline the process, the DNase/RNase treatment reported by Kühner *et al.* [24] was bypassed, as well as treatments with LiCl, EDTA and acetone from older protocols, resulting in only a minimal reduction of the overall fluorescence level, measured as not significant (Fig. S2).

The fluorescent PG was readily employed as a substrate for a fluorescence anisotropy-based assay, with mutanolysin chosen as a benchmark enzyme, given its status as a ‘workhorse’ for HPLC analysis of PG, resulting in a decrease in the anisotropy signal upon addition of the enzyme (Fig. 3, panel A). We noted high background noise during initial assay development (Fig. S3), which required the application of curve smoothing algorithms (GraphPad Prism 10.0). To streamline data processing, and remove any further manipulation, the number of flashes in the ClarioStar reading was increased, resulting in cleaner curves. Regarding this, we found that setting the number of flashes at 50 is a good compromise between background noise and throughput, but especially active enzymes can be analysed with <20 flashes with virtually no effect from background noise. Given anisotropy theoretical independence from sample volume [34], the reaction volume was minimised from 50 µl to 10 µl. A 5 µl reaction volume was also tested and, while the assay itself was not affected, the insoluble nature of the sample alongside evaporation of the aqueous layer, resulted in a sudden and steep increase in the anisotropy signal after ∼85 min (Fig. S4). To overcome this, labelled PG was predigested with mutanolysin and tested for fluorescence against an equally treated unlabelled PG. Monodispersity of the sample was tested via Flow Induced Dispersion Analysis (FIDA, Fig. S5). This last step is important to verify solubility of the substrate, as, while non-solubility does not represent an obstacle [35], under the exact Taylor conditions, dispersion measurements can discriminate between soluble molecules and larger aggregates. The soluble muropeptides were then tested as a substrate in our anisotropy assay using LytA (an amidase from *S. pneumoniae*, not affected by mutanolysin digestion of the PG), revealing that predigestion does not affect the outcome of the assay, rather allowing for a more consistent substrate state (Fig. 3, panel B).

**Figure 2:**
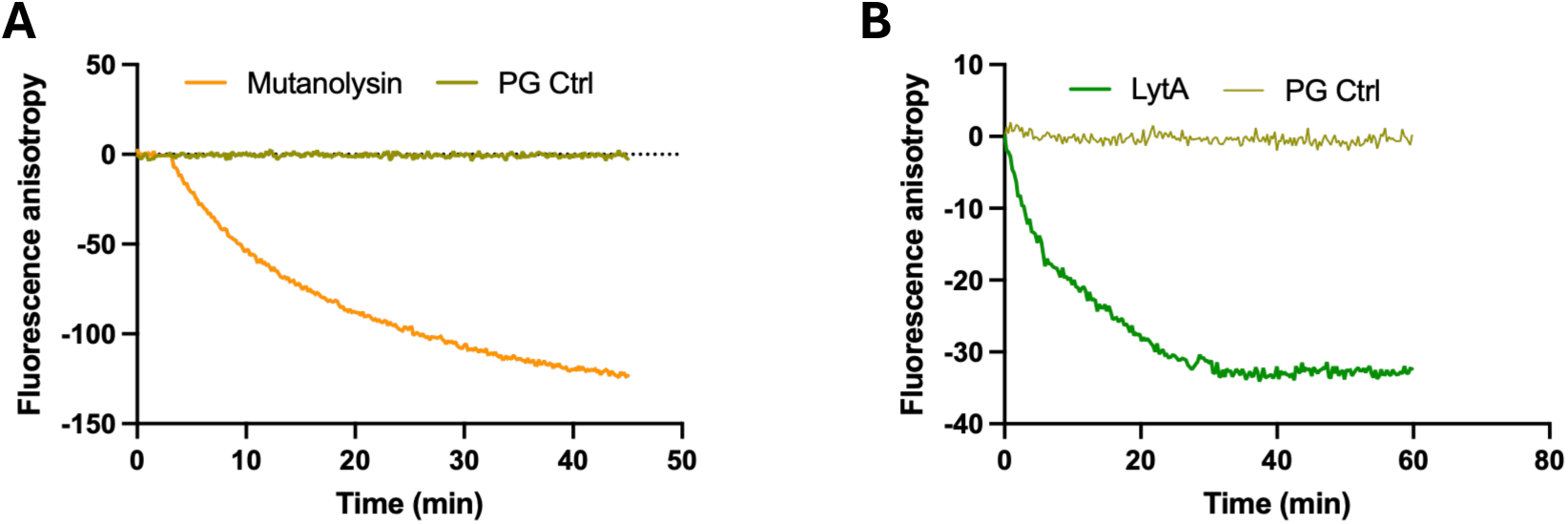
(A) Mutanolysin digestion of pneumococcal PG; the downward curve when the enzyme is added to the reaction mix represents product (GlcNAc-MurNAc-pentapeptide) release upon enzymatic activity. (B) LytA activity on mutanolysin-digested pneumococcal PG, showing activity on the solubilised muropeptides. All figures are representative of one out of three biological repeats.

**Figure 3:**
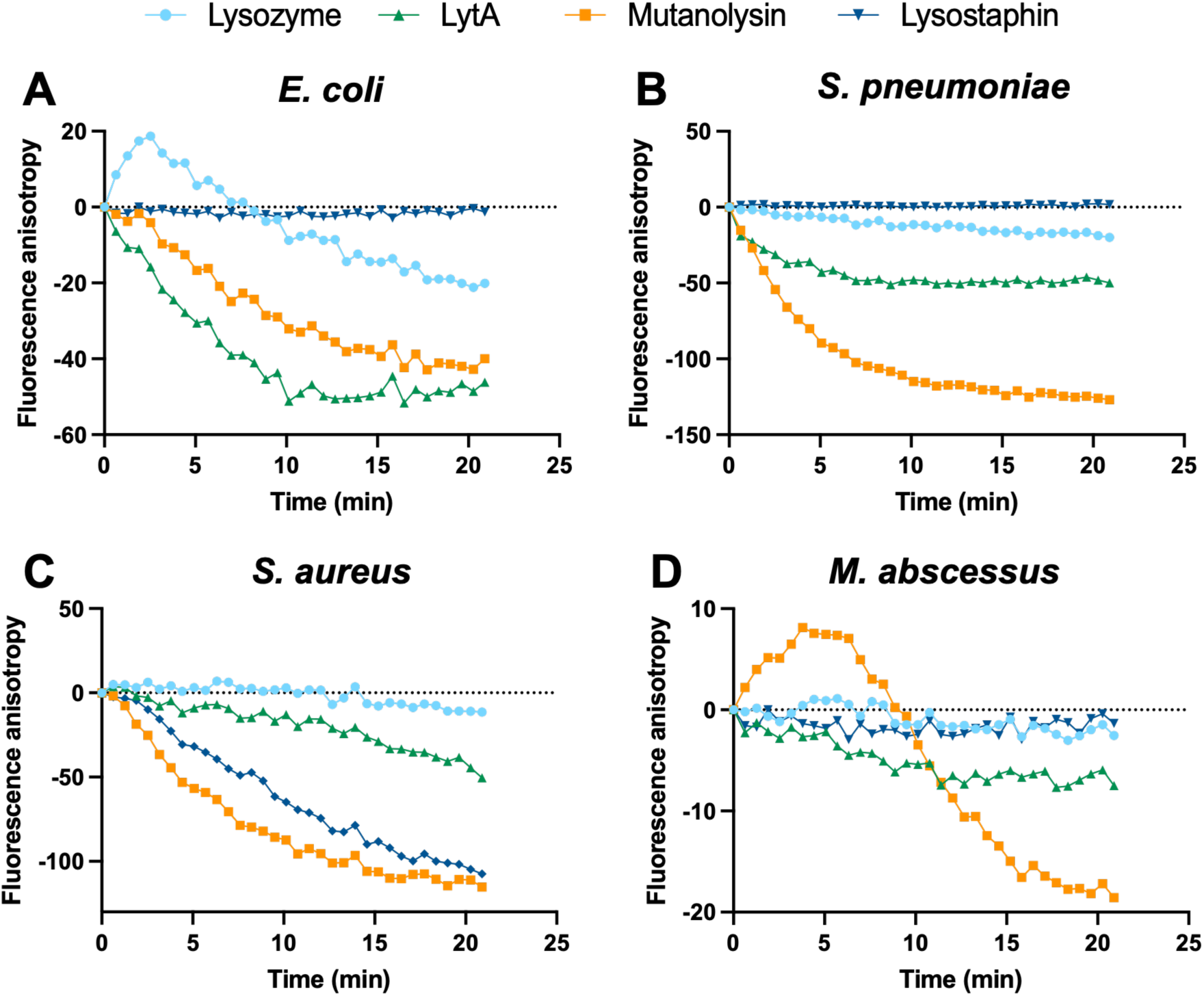
Anisotropy signals from single different substrates assay. (A) *E. coli* PG as substrate, showing activity from all enzymes used; lysozyme shows an increase in the anisotropy signal during the first minutes, possibly due to lower V_max_ compared to mutanolysin, which determines the formation of a substrate-enzyme complex, thus briefly increasing molecule size. (B) *S. aureus* PG as substrate; lysostaphin was employed as substrate-specific enzyme, showing comparable activity to mutanolysin; lysozyme, shows no activity, consistent with *S. aureus* O-acetylation. (C) *M. abscessus* PG as substrate, showing activity from mutanolysin, with a trace shape similar to the one caused by lysozyme with *E. coli*, and partial activity from LytA. (D) *S. pneumoniae* PG as substrate, highlighting activity from both mutanolysin and LytA; lysozyme shows only partial activity, consistent with partial O-acetylation of *Spn* PG. All data are normalised to the same starting position. Blank PG signal was measured and subtracted pairwise to each data point. Each plot is representative of three biological replicates.

### Different substrates and enzymes combinations can be assayed at the same time

One of the major obstacles when working on PG-digesting enzymes is the time-consuming activity assays that have been traditionally employed. The main aim of our anisotropy approach is to deliver a faster, reproducible alternative to the currently used methods for enzyme activity testing. To demonstrate this, we set up a full plate layout experiment, where we tested all combinations of our selected substrates and enzymes (Table S2). PG architectures and enzymes were selected to provide a wider range of substrates and digestion mechanisms in order to test the assay for multiple applications.

The results (Fig. 3) show activities comparable to theoretical expectations for each enzyme on their substrate (Table S2), implying that the assay can be employed to investigate preferred substrates and better understand a reaction mechanism. Specifically, mutanolysin was able to digest all substrates, only showing lower overall activity against *M. abscessus* PG, which may be related to the presence of glycolylated MurNAc saccharide units in mycobacterial PG [45]. The N-glycolyl modification has been shown to negatively impact lysozyme activity [46], and, while mutanolysin activity is not impacted, previous studies failed in measuring real-time substrate degradation. Lysozyme was active against *E. coli* PG, but only partially active against the partially O-acetylated *S. pneumoniae* PG, and completely inactive against the O-acetylated *S. aureus* PG and the N-glycolylated *M. abscessus* [46, 47]. LytA showed activity against all PG substrates, in accordance with its amidase mechanism [43], given the similarity in the *N*-acetylmuramoyl-L-alanine linkage in all employed substrates. Lysostaphin, only showed activity against its preferred substrate, *S. aureus* pentaGly-branched PG.

### Fluorescence anisotropy has potential for high throughput screening of enzyme inhibitors

The main scope of PG-digesting enzymes research is to study those enzymes as potential new targets for the development of effective antimicrobial drugs, given their essentiality in the cell wall machinery, or as synergistic targets to aid increased bacterial sensitivity to known antibiotics. For this reason, a robust activity assay to study PG hydrolases and lyases must offer the possibility of high throughput (HT) screening of inhibitors. This avoids the “double hits” approaches, where an affinity screen is first set up to identify best interactors, followed by traditional activity assays to determine the best inhibitors. To demonstrate our anisotropy-based assay is effective, we determined the IC_50_ of NBS against mutanolysin, using pneumococcal PG as a benchmark substrate. NBS is an organic compound commonly used as a brominating agent in organic synthesis. Whilst NBS is capable of breaking down peptide bonds, this activity is only associated with tryptophyl, tyrosyl and histidyl bonds [48]. An initial trial was set up to determine if the assay could detect inhibition (Fig. 5, panel A), showing a flatlined anisotropy signal in presence of 1 mM NBS, a concentration previously reported to inhibit mutanolysin activity [22]. Possible reaction between NBS and PG was tested and deemed not significant (Figure S6).

We then set up a 3-fold titration of NBS starting at 3 mM to determine its IC_50_ (Fig. 5, panel B, Fig. S6), fitting the result to 25.16 ± 0.08 µM. This inhibition test also allowed for a more robust Z’-score calculation, to provide a mathematical justification for the use of our anisotropy-based assay in the context of HT screening [49, 50], resulting in a calculated Z’-score of 0.69, with values above 0.5 being considered as the minimum to achieve HT [51, 52].

**Figure 4:**
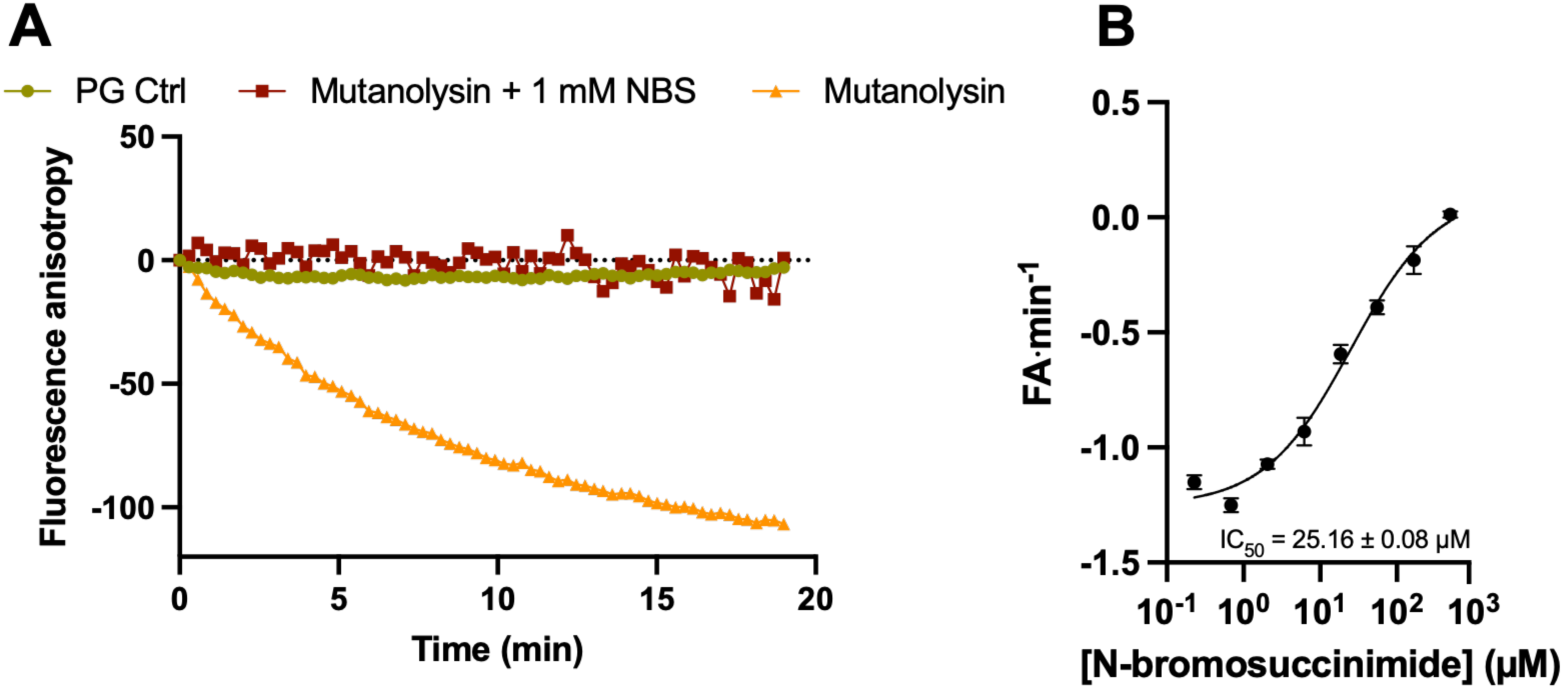
(A) Proof of concept for mutanolysin inhibition by N-bromosuccinimide, showing clear muramidase activity by mutanolysin alongside non-detectable activity from mutanolysin preincubated with NBS. (B) IC_50_ fitting of NBS on mutanolysin, calculated at 25.16 ± 0.08 µM; IC_50_ calculation was carried out via Inhibitor vs. response nonlinear fitting analysis on GraphPad Prism 10.0, with R^2^ = 0.9842. The upward direction of the curve is due to the nature of the anisotropy assay, where an active enzyme will generate a downward curve, and so a negative initial velocity value. All graphs are representative of one out of three biological replicates.

### Limitations of the assay

Whilst proven effective in measuring activity and inhibition for PG-digesting enzymes, the assay described in this study is not free of limitations. The insoluble nature of PG makes it a difficult substrate to extrapolate accurate kinetics, as it presents poor monodispersity, thus losing consistency between wells. In our attempt, we used a general quantification based on dry weight of PG and, while an approximation of enzyme kinetics can be made this way (Fig. S7), it does not represent an accurate and robust measurement. Further optimisation, possibly based on FIDA-derived molecular weight of mutanolysin-digested PG, could retrieve more accurate measurements for amidase and endopeptidase classes (not impacted from modifications of the glycan strand). Muramidases and glycosidases would require a different approach, as their preferred substrate architecture would be lost with digestion.

A second limitation is due to the intrinsic inability of the assay to retrieve information about the structure of the product, making it prohibitive to use it for enzyme classification. Mass spectrometry remains the gold standard for that application.

## CONCLUSIONS

Bacterial PG is a polymeric structure surrounding and protecting the cell, with shape-determining and scaffolding roles that are stably intertwined with essential protein supercomplexes dedicated to its remodelling during growth and cell division [1, 10, 11]. The essentiality of processes involving PG have been the target for generations of antibiotics, and antimicrobial discovery still heavily relies on targeting cell wall-related enzymes [16]. In this context, PG-digesting enzymes, responsible for facilitating septal separation [53] or in some cases working as modulators of immune activation [14], represent a class of inviting targets to disrupt the division process. As for all enzymes, inhibitor discovery requires robust methods to screen for libraries of possible interactors, a task made more complicated by the fact that PG degradation analysis often relies on suboptimal turbidity screenings [21, 22]. Turbidity measurements also often translate to low throughput, time- and resource-consuming methods, thus only sufficient for compound finalisation. Some cysteine peptidases, like CHAP and NlpC/P60 domain-containing proteins [16], can be probed with sulfonyl compounds releasing a fluorescent molecule upon formation of a disulfide bond with the active free Cys. This allows to avoid turbidimetry and enables the identification of compounds that are covalently interacting with the active site [54], but does not represent an actual activity assay.

With the aim of offering a faster, standardisable and reliable method with high throughput capabilities, we used fluorescence anisotropy, a technique already employed in multiple molecular interaction analysis settings, as well as in enzymology, and recognised by regulatory bodies as a robust tool for drug discovery, to detect digestion of labelled PG. To this end, we were able to isolate the labelled PG following an optimised protocol whilst still retaining its fluorescence and employ it in an easy to set up anisotropy assay with known PG-targeting enzymes, both commercially available and purified in house. Our assay can at the same time be used to screen for inhibitors, showing HT potential, as proven by the N-bromosuccinimide effect on mutanolysin. The volume miniaturisation, alongside the ease of sample preparation, allow for the analysis of various combinations of substrates and enzymes, which we believe is the strongest feature of this novel application of fluorescence anisotropy.

Given the general fluorescence anisotropy principle [26], combined with the polymeric nature of PG, our method can potentially be applied to any fluorescently labelled polymer, with the only limiting factor being the labelling process itself.

The final product is a fast, reliable and cost-effective assay, which we speculate will aid in finding novel inhibitors for difficult to analyse enzymes.

## AUTHOR INFORMATION

### Authors contribution

S.C. devised and developed the assay, performed PG extraction from *S. aureus*, *S. pneumoniae* and *E. coli* and designed the FIDA analysis; C.M.T. extracted *M. abscessus* PG and performed FIDA analysis; N.S.B. cloned and purified LytA. D.I.R. supervised the investigation. S.C. and D.I.R. wrote the initial draft, and all authors revised and approved the manuscript.

## ACKNOWLEDGMENTS

The authors thank Prof. Malcolm Winkler from the Indiana University, Bloomington for the kind donation of *S. pneumoniae* strain D39 in its WT and capsule deficient (Δ*cps*) variants.

The Authors acknowledge use of the FIDAneo instrument provided by the Bio-Analytical Shared Resource Laboratories within the School of Life Sciences, University of Warwick.

