## Supplementary data for "Development of a high throughput continuous fluorescence anisotropy-based assay to analyse peptidoglycan-digesting enzymes"

| LT1 | ATGGAAATTAATGTGAGTAAATTAAGAACA |
| --- | --- |
| LT2 | TTATTTTACTGTAATCAAGCCATCTGGCTC |
| LTG1 | CACAGCAGCGGCCTGGTGCCGCGCGGCAGCATGGAAATTAATGTGAGTAAATTAAGAACA |
| LTG2 | TTCCTTTCGGGCTTTGTTAGCAGCCGGATCTTATTTTACTGTAATCAAGCCATCTGGCTC |

**Table S1:** primers used to clone the LytA construct into a pET-28b(+) vector.

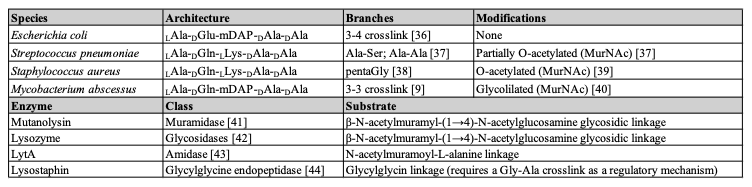

**Table S2:** various peptidoglycan architectures used in the study; enzyme classes and specific substrates assayed via fluorescence anisotropy.

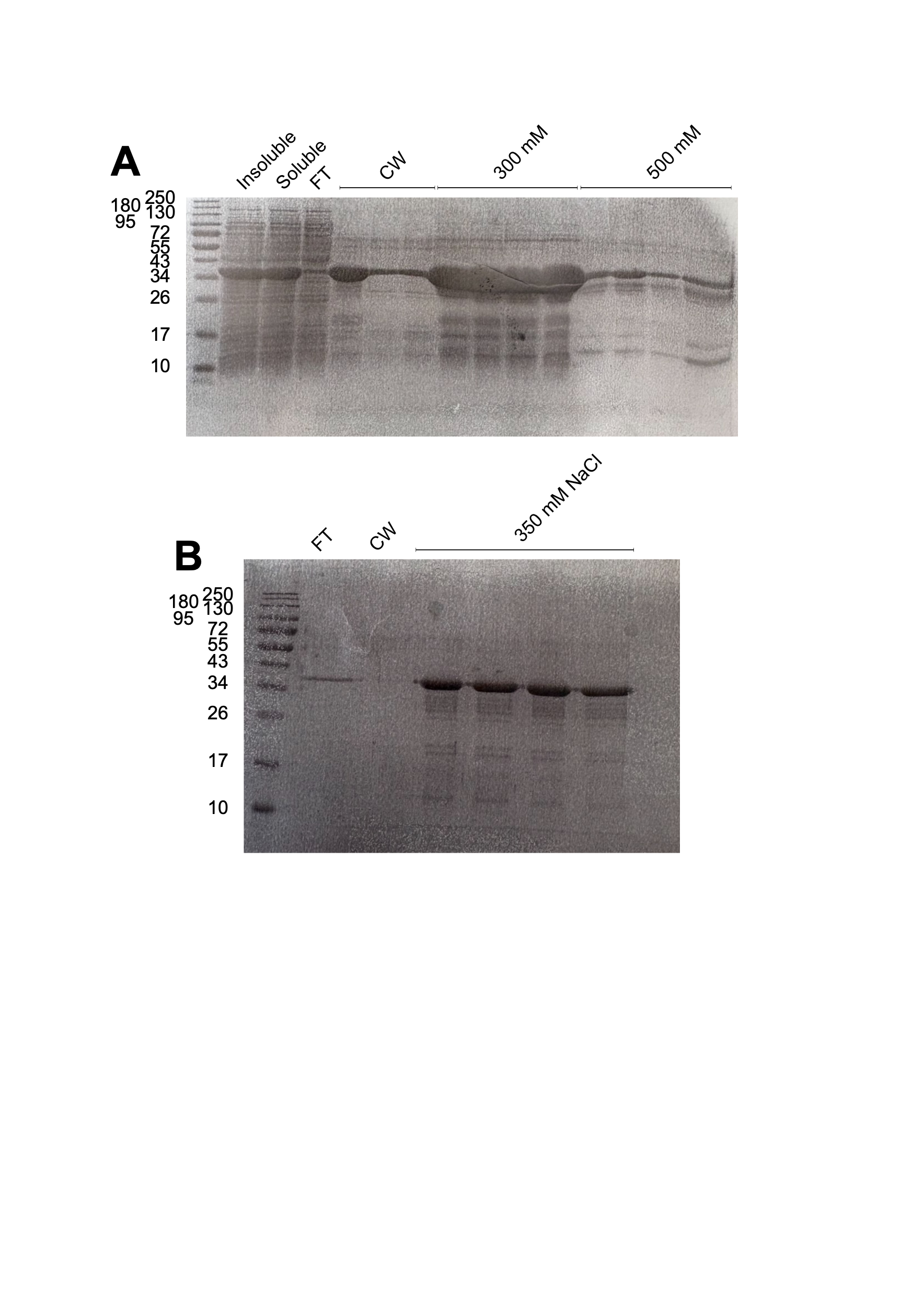

**Figure S1:** A) IMAC SDS-PAGE. FT = flowthrough; CW = column wash; imidazole concentration indicated as mM on legend. B) Anion exchange SDS-PAGE. Samples to be loaded were selected after analysing the absorbance chromatogram at λ = 280 nm.

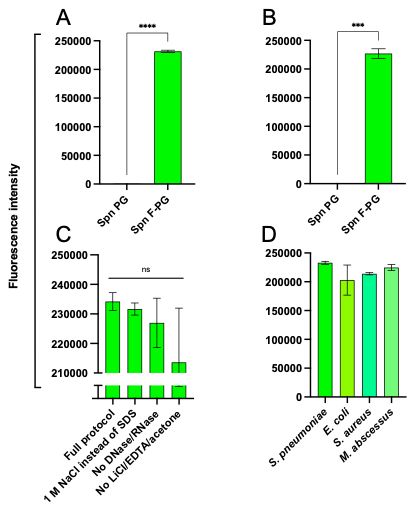

**Figure S2:** Fluorescence intensity of extracted PG. A) Fluorescence intensity after standard extraction. B) Fluorescence intensity after large scale (2 L) extraction. C) Intensity comparison for different species PG used in this study after standard extraction. D) Intensity comparison for different variation of the extraction protocol, showing no significant difference.

Statistical evaluation carried via Student *t* test (GraphPad Prism 10.0).

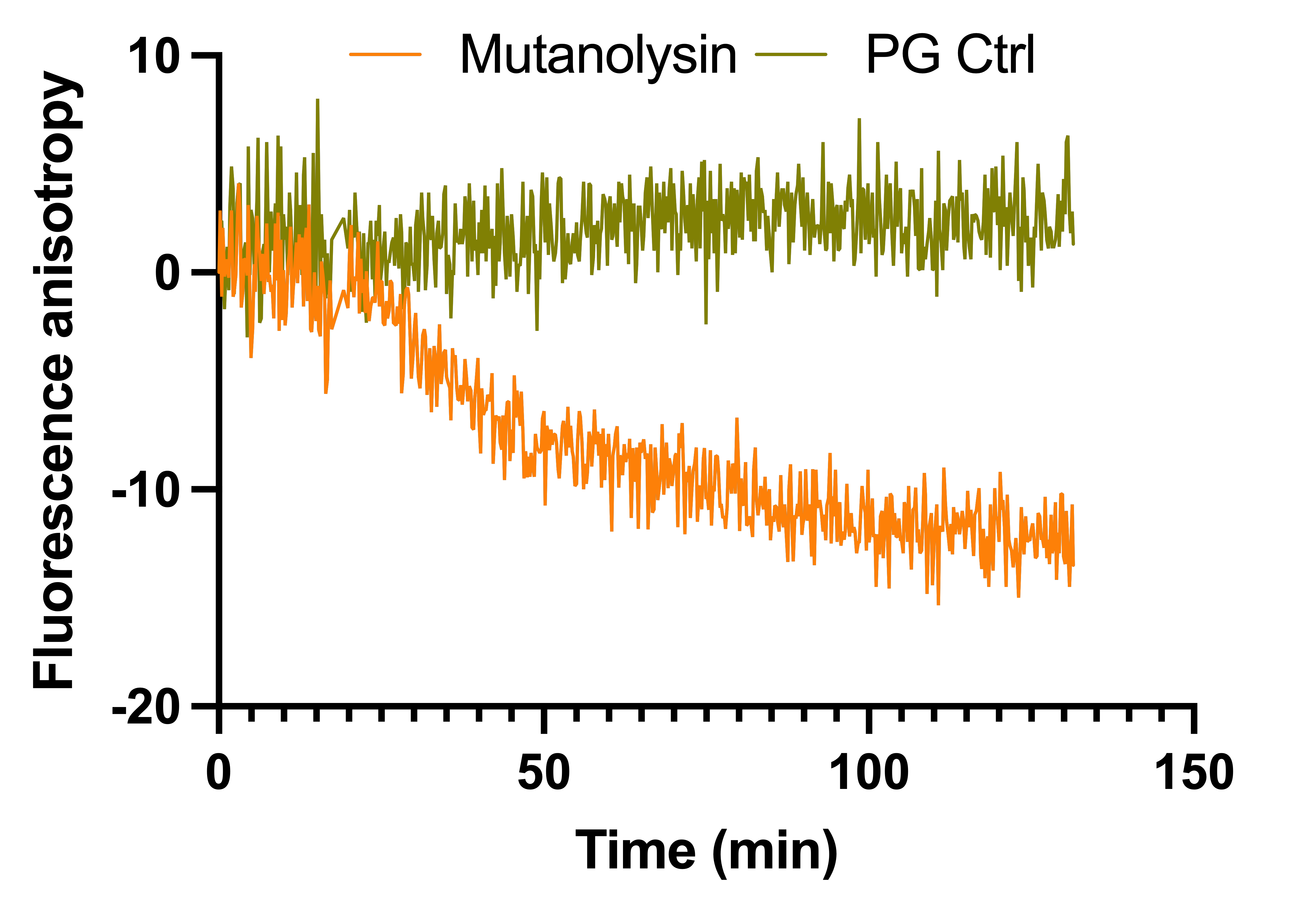

**Figure S3:** Proof of concept analysis of mutanolysin activity on pneumococcal PG, showing clear activity with high background noise

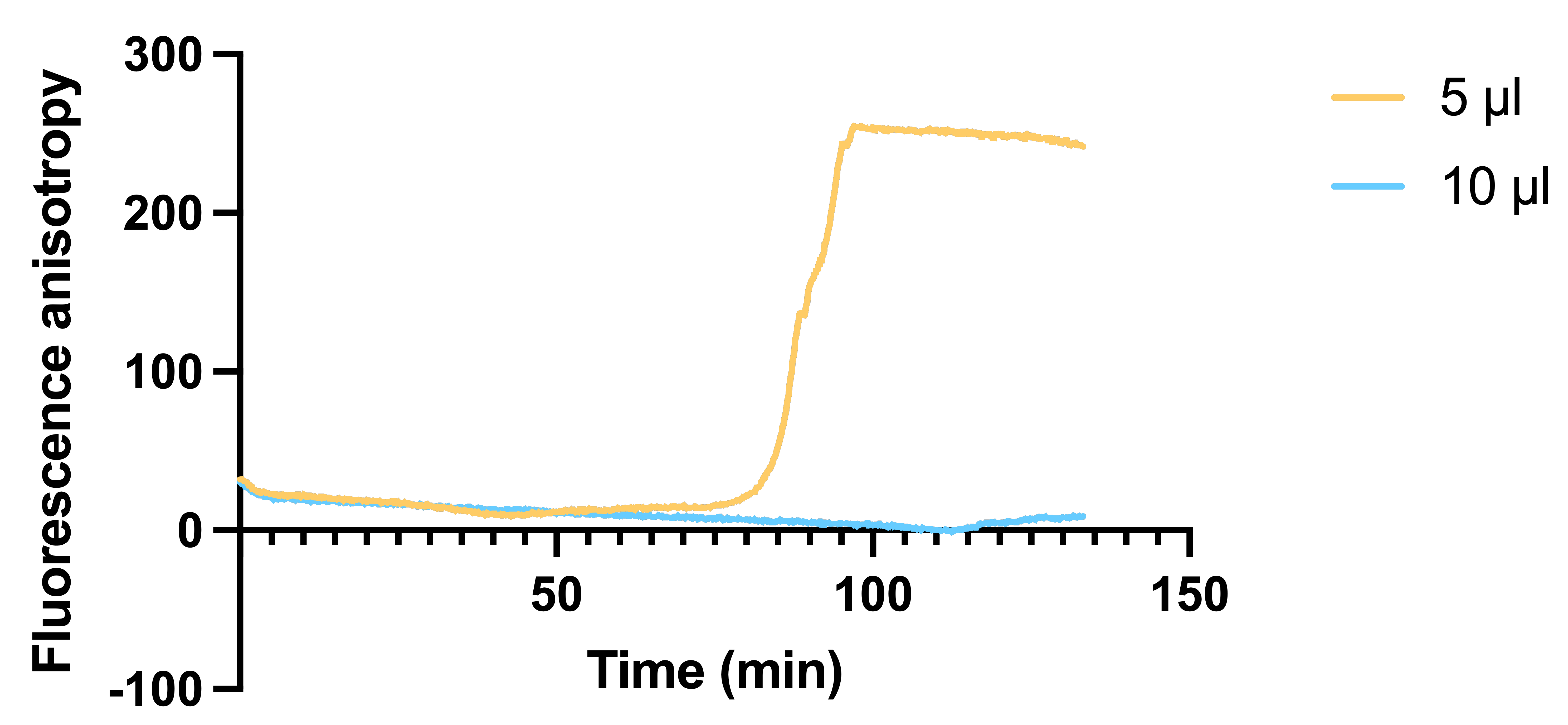

**Figure S4:** Volume miniaturisation test. Only substrate was used in this experiment, resulting in significant increase in anisotropy, possibly due to buffer evaporation, in the 5 µl wells after ~85 min. The 10 µl wells only show minimal increase in the anisotropy signal at the 2 h mark.

**Figure S5:** Substrate titration experiment. A) Fluorescence anisotropy traces. B) Michaelis-Menten fitting of initial velocities calculated form the anisotropy traces. C) Fitting results.

Fitting calculations were carried on GraphPad Prism 10.0. The results shown are representative of one out of three biological replicates.

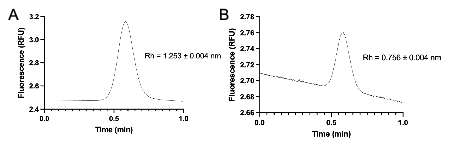

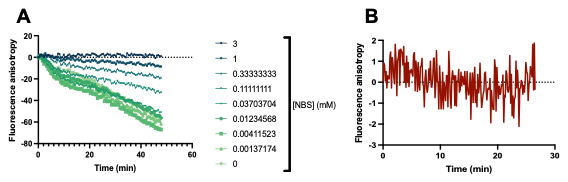

**Figure S6:** A) Fluorescence anisotropy trace for N-bromosuccinimide inhibition of mutanolysin. B) Anisotropy signal of 3 mM NBS added to S. pneumoniae PG, showing no detectable reaction.

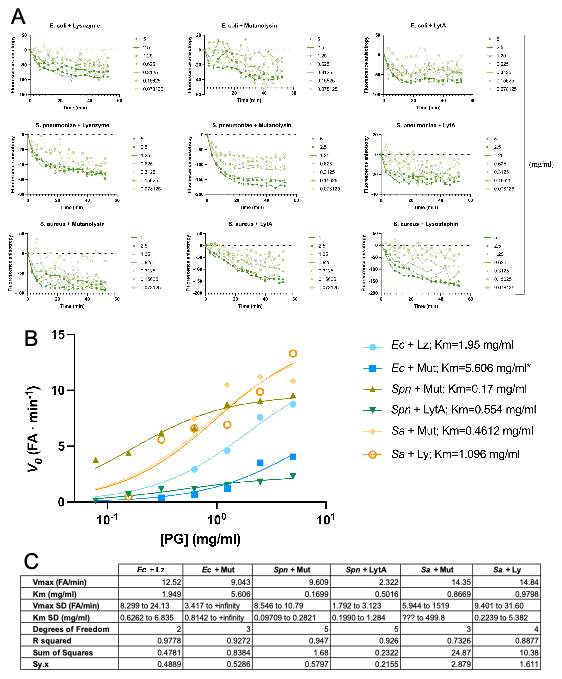

**Figure S7:** Flow Induced Dispersion Analysis of mutanolysin-digested peptidoglycan, showing a difference in hydrodynamic radius (Rh) consistent with PG branching and crosslinking of the different species. FIDA analysis shows no sign of aggregares in the indicator molecule, indicating a monodisperse and fully soluble sample. A) S. pneumoniae PG. B) E. coli PG.
